# *Gnas*^R201C^ Induces Murine Pancreatic Cystic Neoplasms through Suppression of YAP1 Signaling and Transcriptional Reprogramming

**DOI:** 10.1101/310292

**Authors:** Noboru Ideno, Hiroshi Yamaguchi, Bidyut Ghosh, Sonal Gupta, Takashi Okumura, Catherine G Fisher, Laura D Wood, Aatur D. Singhi, Masafumi Nakamura, J Silvio Gutkind, Anirban Maitra

**Affiliations:** Department of Translational Molecular Pathology Houston, Texas 77030, USA; Sheikh Ahmed Center for Pancreatic Cancer Center Houston, Texas 77030, USA; Department of Pathology Baltimore 21287, USA; Department of Oncology Sol Goldman Pancreatic Cancer Research Center Baltimore 21287, USA; Department of Anatomic Pathology, University of Pittsburgh, Pittsburgh 15260, USA; Department of Surgery and Oncology, Graduate School of Medical Sciences, Kyushu University, Fukuoka, Japan; Department of Pharmacology, University of California San Diego, La Jolla, California 92093, USA

**Author notes:** **Correspondence:** Noboru Ideno, MD, PhD, Department of Translational Molecular Pathology, Sheikh Ahmed Center for Pancreatic Cancer Research, The University of Texas MD Anderson Cancer Center, Houston, Texas 77030, USA.

**Keywords:** Intraductal papillary mucinous neoplasm, IPMN, *GNAS*, Hippo pathway, YAP1

## Abstract

**Background & Aims:** Somatic “hotspot” mutations of *GNAS*, which encodes for the alpha subunit of stimulatory G-protein, are present in ~60% of intraductal papillary mucinous neoplasms (IPMNs) of the pancreas. There are currently no cognate animal models that recapitulate the biology of mutant *Gnas*-induced IPMNs, and the underlying mechanisms that lead to the cystic pathway of neoplasia in the pancreas remain unknown.

**Methods:** We generated *p48*-Cre; LSL-*Kras*^G12D^; Rosa26R-LSL-rtTA-TetO-*Gnas*^R201C^ mice (*Kras*; *Gnas* mice) where pancreas-specific *Gnas*^R201C^ expression was induced by doxycycline administration. In this model, mutant *Kras* is constitutively expressed, and control mice were produced through absence of doxycycline. Separate cohorts of mice were utilized for timed necropsies and for Kaplan-Meier survival analysis. Isogenic cell lines (with doxycycline inducible mutant *Gnas* expression) were propagated from the resulting pancreatic ductal adenocarcinoma (PDAC).

**Results:** Co-expression of *Kras*^G12D^ and *Gnas*^R201C^ resulted in the development of pancreatic cystic lesions resembling human IPMNs in 100% of mice, with higher grades of epithelial dysplasia observed over time. Approximately one-third of *Kras*; *Gnas* mice developed PDAC at a median of 38 weeks post doxycycline induction. *Gnas*^R201C^ did not accelerate oncogenic transformation with *Kras*^G12D^, but rather, reprogrammed Ras-induced neoplasms towards a well-differentiated phenotype. *Gnas*^R201C^ induction led to activation of the inhibitory Hippo kinase cascade and cytoplasmic sequestration of phosphorylated YAP1 protein, a phenomenon that was also observed in human IPMN with *GNAS* mutations.

**Conclusions:** *GNAS*^R201C^ functions not as a traditional oncogene, but rather as an “oncomodulator” of *KRAS*-mediated pancreatic neoplasia, through suppression of YAP1 and transcriptional reprogramming towards a differentiated (large ductal) phenotype.

## INTRODUCTION

Pancreatic ductal adenocarcinoma (PDAC) is the 3^rd^ most common cause of cancer-related mortality in the United States ^1^. The multistep progression of PDAC can occur on the backdrop of non-cystic, microscopic precursor lesions known as pancreatic intraepithelial neoplasias (PanINs), or alternatively, through a cystic pathway that portends a distinct natural history ^2^. Overall, pancreatic cysts are identified in greater than 2 % of the general patient population, rising to 10% amongst individuals above 70 years of age ^3^. Many of the cystic lesions are essentially benign, and harbor no risk of progression to PDAC. In contrast, mucin-secreting cysts of the pancreas, specifically intraductal papillary mucinous neoplasms (IPMNs), are *bona fide* precursor lesions of PDAC ^3^. Patients with non-invasive IPMNs have a 5-year survival rate of 90–100 % upon surgical resection, which is reduced to about 50% for those with an invasive component ^4^. Nonetheless, in contrast to PDAC that arises through the non-cystic (PanIN) pathway to invasive neoplasia, and which is nearly always fatal, cancers arising in the context of IPMNs demonstrate a more indolent course. Further, even within the universe of IPMNs, the vast majority does not progress to PDAC, although the biological basis for this favorable outcome has, thus far, remained elusive.

Progress in translational research on pancreatic cystic lesions has been hampered by lack of animal models that recapitulate the biology and genetic of the cognate human disease. In 2011, we performed a comprehensive next generation sequencing (NGS) assessment of IPMNs, and identified two recurrent driver alterations, oncogenic mutations of *KRAS* and *GNAS* ^5^. Specifically, activating mutations of *KRAS* were observed in 80% of cases, while activating mutations of *GNAS*, which encodes for the alpha subunit of a stimulatory G-protein (Gαs protein), were present in 66% of IPMNs. *GNAS* mutations were restricted to codon 201, a previously described “hotspot” in epithelial neoplasms. Overall, 96 % of IPMNs harbored either *KRAS* or *GNAS* mutations ^5, 6^, underscoring the importance of these alternations in IPMN pathogenesis. Further, our prior studies have confirmed that *GNAS* mutations are observed early in IPMN pathogenesis, including in precursor lesions of IPMNs, the so-called “incipient IPMNs”, establishing these mutations as an early driver of mucinous neoplasms of the pancreas ^7^. However, the precise functional role played by mutant Gαs protein in cooperation with oncogenic Ras during IPMN progression has been incomplete due to lack of an autochthonous model that recapitulates the cognate human disease process.

Under homeostatic conditions, Gαs transiently activates adenylate cyclase and increases cyclic adenosine monophosphate (cAMP) production as a second messenger. Although Gαs possesses intrinsic enzymatic activity that hydrolyzes bound GTP to GDP, thereby returning them to active form, *GNAS* R201 mutations markedly reduce efficiency of GTP hydrolysis that render Gαs constitutively active. This results in sustained elevation of the intracellular cAMP level ^8^, which subsequently activates protein kinase A (PKA) and downstream signaling cascades ^9^. Activating mutations of *GNAS* were first described in the context of endocrine neoplasms like pituitary adenomas and juvenile granulosa cell tumors ^8, 10, 11^, and subsequently, in epithelial tumors, such as in a minor subset of colorectal cancers ^12^. The recent generation of a genetically engineered mouse (GEM) model of conditional (doxycycline inducible) *Gnas*^R201C^ activation by Gutkind and colleagues ^13^ has now allowed us to examine the role of constitutively active Gαs in IPMN pathogenesis. While our GEM model faithfully recapitulates the multistep progression of human IPMNs, we also elucidate the molecular mechanisms through which Gαs elaborates a cystic differentiation with the context of mutant Ras signaling. Specifically, we demonstrate that Gαs does not behave as a traditional cooperating oncogene, but rather as a modulator of differentiation of Ras-induced pancreatic neoplasia (“*onco-modulato*”) through attenuation of YAP1 signaling within epithelial cells. Our studies likely clarify the natural history of IPMNs and the low rate of progression to invasive neoplasia on the background of cystic precursors.

## METHODS

### Genetically engineered mice

All animal studies were carried out according to the MD Anderson Institutional Care and Use of Animals Committee (IACUC)-approved protocols. To model the co-expression of mutant *Kras* and *Gnas* in the murine pancreatic epithelium, we used a previously described doxycycline (doxy)-inducible *Gnas*^R201C^ model ^13^. In this model, tissue-specific expression of *Gnas*^R201C^ is regulated by crossing the mice to an appropriate *Cre*-driver (in this case, p48-*Cre*). This leads to removal of a lox-STOP-lox (LSL) cassette upstream of a reverse tetracycline transactivator (rtTA) and expression of rtTA in the pancreatic epithelium. In the presence of doxy, rtTA binds to the Tet operon (TetO) upstream of *Gnas*^R201C^ leading to transgene expression in the pancreas ^13, 14^ We generated compound heterozygous *p48*-Cre; LSL-*Kras*^G12D^; Rosa26R-LSL-rtTA-TetO-*Gnas*^R201C^ mice (*Kras*; *Gnas* mice) (**Figure 1A**). Doxycycline diet was fed at a dose of 0.0060 %. The time course for supplementation is outlined in the **Figure 1B**. The confirmation of recombination of *Kras*^G12D^ and transcription of *Gnas*^R201C^ are shown in the pancreatic lysates in **Supplementary Figure 1A and B**.

**Figure 1.**
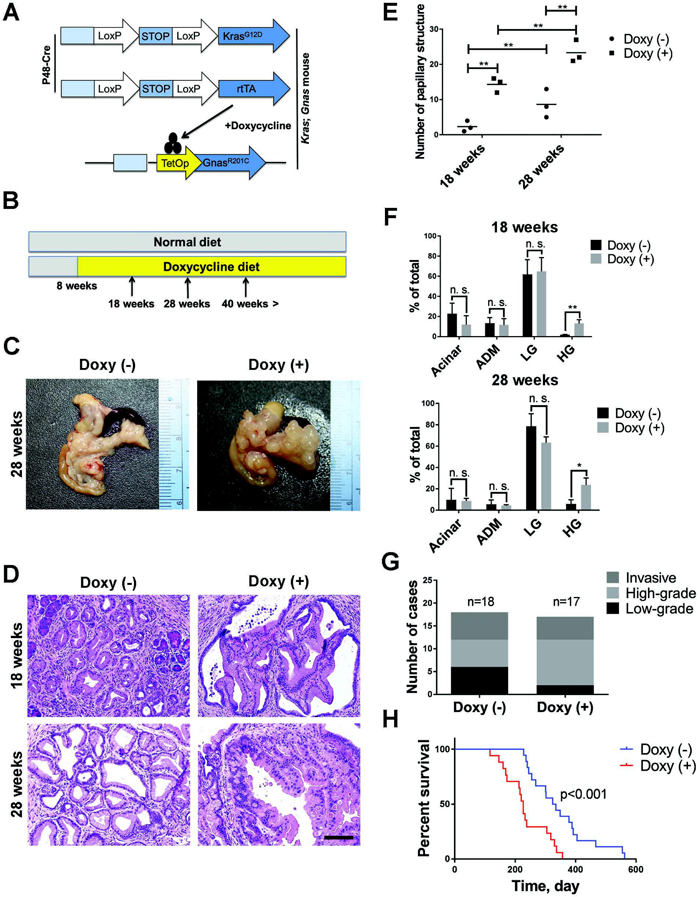
Concomitant expression of *Gnas*^R201C^ and *Kras*^G12D^ mutations in the murine adult pancreas induces the development of cystic neoplasms resembling human IPMN. (A) Schema for conditional activation of pancreas-specific *Gnas*^R201C^ expression using doxycycline (doxy) on a background of constitutive *Kras*^G12D^ expression. (B) *Kras*; *Gnas* mice were maintained on a doxy diet from the age of 8 weeks to induce *Gnas*^R201C^. Littermate *Kras;Gnas* mice without doxy served as isogenic controls. (C) Gross pathology of the pancreas in *Kras;Gnas* mice maintained without doxycycline (left) or with doxycycline (right), showing multiple cystic lesions throughout the pancreas of latter mice (time of harvest = 28 weeks). (D) Representative pictures of pancreatic precursor lesions arising in mice with or without doxy induction, including murine pancreatic intraepithelial neoplasia (mPanIN) in control mice pancreata and intraductal papillary lesions in *Kras*; *Gnas* mice fed doxy diet until the indicated time points (scale bar, 50μm). (E) Quantification of papillary structures in pancreata with or without *Gnas*^R201C^ expression at the indicated time points. (F) Quantification of pancreatic lesions in *Kras*; *Gnas* mice pancreata with or without *Gnas*^R201C^ expression at the indicated time point. (G) Quantification of the disease stages and histology in *Kras*; *Gnas* mice without doxy (n=18) and with doxy administration (n=17). (H) Kaplan-Meier survival analysis for *Kras*; *Gnas* mice without doxy (n=18) and with doxy administration (n=17).

### Histological evaluation of pancreas

To histologically evaluate the multistep progression of pre-invasive neoplasms, including mPanINs or murine IPMNs (mIPMNs), the grade of ductal lesions was evaluated in representative pancreatic sections of cohorts of *Kras*; *Gnas* mice, with or without doxycycline at the age of 18, 28, and more than 40 weeks, respectively (**Figure 1B**). Only the highest-grade lesion per pancreatic lobule was evaluated, using published criteria for low-grade or high-grade mPanIN and mIPMN ^15–17^ An average of 171 total pancreatic lobules were counted from 3 independent animals for each group. The data were expressed as a percentage of the total counted pancreatic lobules. Cystic papillary lesions in our mouse model were defined as epithelium-lined cystic structures >500µm, and composed of papillary epithelial proliferations with fibrovascular cores and varying degrees of cellular atypia, which is a *sine qua non* of human IPMNs ^15^. Cystic papillary structures in the representative pancreatic sections which met the definition were counted to assess the neoplastic nature of *Kras*; *Gnas* mice. In terms of invasive carcinomas, they were graded based on the most poorly differentiated component when multiple histopathlogical grades were observed.

### Establishment of primary pancreatic ductal adenocarcinoma cell lines with doxycycline inducible mutant *Gnas*^R201C^

Freshly isolated murine PDAC tissues were minced by sterile scalpel and washed with PBS containing 1% penicillin and streptomycin and re-suspended in RPMI with 10 % fetal bovine serum (FBS) and 1% penicillin and streptomycin, and seeded in 60 mm collagen coated dishes. Five days after seeding of tumor pieces, residual tissue was removed and attached colonies comprised of murine PDAC cells were isolated using 150µl cloning cylinders (Sigma-Aldrich, St Louis, MO) and expanded. For the purpose of this study, the two cell lines used in all of the experiments are designated as “LGKC-1” and “LGKC-2”, respectively. Both cell lines were used at less than passage 10 to minimize passage-dependent artifacts ^18^.

### Gene expression analysis and reverse-transcription quantitative polymerase chain reaction

Total RNA extraction from cultured LGKC-1 and LGKC-2 cells, as described previously ^19^. Microarray analysis was performed with the Agilent Mouse GE 4×44k v2 Microarray Kit (Agilent, Santa Clara, CA) using mRNA from mPDAC cell lines established from primary PDAC in the *Kras*; *Gnas* mice. These cells express mutant *Kras* constitutively, while mutant *Gnas* expression can be turned on or off by doxycycline administration, producing and isogenic system. To identify differentially enriched gene sets, the raw intensities were quantile normalized and the data were subjected into Gene set enrichment analysis (GSEA) software, v2.2.3 ^20, 21^. Gene sets collection from oncogenic signatures gene sets (v5.2) were used in the analysis. For RT-qPCR protocol, see **Supplemental Materials and Methods**.

### Colony formation and invasion assays

Anchorage-independent growth in soft agar was assessed as described previously ^22^. Invasion assay was performed as described previously ^19^.

### Allograft model for assessing tumor growth rates and differentiation

For *in vivo* growth and differentiation assays, murine PDAC cells (with or without additional genetic modifications of *mYap1* as described in the text) were suspended in 100µl of 50% Matrigel/PBS mix and injected subcutaneously into the lower flanks of age-matched female NUDE mice (n=4 per group). Twenty four hours after injection, the mice were randomized to be fed normal diet or doxycycline diet (0.6%). Tumor volumes were measured every 3-4 days for 17 days and tumor weight was measured at the end of the experiment. Histological assessment was conducted for grade of tumor differentiation, and additional immunohistochemistry and western blot analyses on the explanted tumors were conducted, as described below.

### Immunofluorescence and Immunohistochemistry

For immunofluorescence, LGKC cells were seeded on the collagen coated glass bottom dishes (Thermo fisher scientific, Grand Island, NY) and incubated 48 hours with RPMI media containing 10% FBS. Cells were incubated for 24 hours in the media with either PBS or 1µg/ml doxycycline to induce mutant *Gnas* and fixed with 100% cold methanol. The cells were blocked in 1xPBS, 1% BSA and 0.3% Triton X-100 for 1 hour followed by incubation with primary antibodies listed in **Supplementary Table 1** at 4ºC overnight. After washing, cells were incubated in conjugated secondary antibody. The proportion of cells per field showing YAP1 localization in the nucleus or cytoplasm was quantified. In 3 independent experiments, 4 fields with 30 to 50 cells were counted per condition per experiment. For immunofluorescence of paraffin-embedded samples, tissue sections were deparaffinized with xylene and hydrated by graded series of ethanol washes. Antigen retrieval was accomplished by heating the slides in citrate buffer at 100ºC for 12 minutes. Nonspecific binding was blocked with serum free protein block (Dako, Glostrup, Denmark) before overnight incubation with primary antibodies listed in the Supplemental Table 1. Sections were incubated with conjugated secondary antibody. Immunohistochemstry on archival human IPMNs samples was performed as described previously ^19^.

### Protein extraction and Western blotting

Protein extraction and western blotting were performed as described previously ^19^. Additional details on methodology and assessment are provided in the **Supplementary Materials and Methods**.

### Statistical analysis

All analyses were performed in triplicate or greater and the means obtained were used for independent t-tests. Statistical analyses were carried out using the Prism 6 statistical analysis program (GraphPad Software, La Jolla, CA). Asterisks denote statistical significance (nonsignificant or NS, P>0.05; *P<0.05; **P<0.01; and***P<0.001). All data are reported as mean ± standard error of the mean (s. e. m.).

## RESULTS

### Expression of *Gnas*^R201C^ in adult *p48-Cre*; LSL-Kras^G12D^; Rosa26R-LSL-rtTA-TetO-Gnas^R201C^ (*Kras*; *Gnas*) mice leads to cystic precursor lesions resembling human IPMN

To determine how *Gnas* mutations alter the natural history of *Kras*-driven murine pancreatic neoplasia, we fed doxycycline (doxy) diet to adult *Kras*; *Gnas* mice starting at the age of 8 weeks to induce *Gnas*^R201C^ on a constitutive mutant *Kras*^G12D^ expressing background (**Figure 1A, B**). In the absence of doxy, i. e. with mutant *Kras*^G12D^ expression alone, the pancreata mainly demonstrated murine pancreatic intraepithelial neoplasias (mPanINs), which are non-cystic precursor lesions of PDAC (**Figure 1C, left**). However, upon co-expression of mutant *Gnas*^R201C^, 100 % of the mice developed cystic lesions beginning as early as 10 weeks of doxy treatment (**Figure 1C, right**). The neoplastic cells expressed apomucins Mucin1 and Mucin5ac but not Mucin2, which resembles the phenotype pancreatbiliary subtype of human IPMNs (**Supplemental Figure 2A**).

To further evaluate the nature of pancreatic precursor lesions with or without mutant *Gnas*^R201C^, the number of papillary structures with fibrovascular cores, which are a diagnostic *sine qua non* of IPMNs, was counted in representative pancreatic sections from doxy administered *Kras*; *Gnas* mice at the ages of 18 and 28 weeks, respectively. For each comparison, littermate *Kras*; *Gnas* mice with no doxy (i. e. with mutant *Kras*^G12D^ expression alone) were used as control. The number of papillary structures was significantly increased in the pancreata of mice co-expressing mutant *Kras*^G12D^ and *Gnas*^R201C^, in comparison to those with mutant *Kras*^G12D^ at both time points. In addition, more papillary structures were observed in the pancreata of the mice at the later time point, suggesting co-expression of mutant *Gnas*^R201C^ progressively enhances the IPMN-specific phenotype (**Figure 1D, E**).

The distribution for grades of all exocrine pancreatic precursor lesions including acinar-ductal metaplasia (ADM), mPanIN, and murine IPMN (mIPMN) were also assessed to determine the effect of mutant *Gnas*^R201C^ on multistep progression of *Kras*^G12D^-driven neoplasia (**Figure 1F**). The proportion of pancreatic parenchyma affected by ADM or low-grade mPanIN or mIPMN was not significantly different, irrespective of the co-expression of mutant *Gnas*^R201C^ at the indicated time points of 18 and 28 weeks (**Figure 1F**). However, in terms of high-grade lesions, the percentage of pancreatic lobules with high-grade (HG) lesions were significantly higher in the pancreata with mutant *Kras*^G12D^ and *Gnas*^R201C^ co-expression than those in the pancreata with mutant *Kras*^G12D^ alone (1.9 v. s. 13.1 %, p=0.006, and 5.8 v. s. 23.6 %, p= 0.016 in 18 and 28 weeks old mice, respectively, **Figure 1F**). These results indicate that *Gnas*^R201C^ co-expression accelerates formation of precursor lesions in the setting of mutant *Kras*^G12D^.

In spontaneously aged mice to 40 weeks and beyond, irrespective of the administration of doxy, virtually the entire pancreatic parenchyma was eventually replaced by ductal neoplastic lesions intermingled fibrotic areas due, in part to the constitutive expression of the mutant *Kras*^G12D^ allele. Overall, 10 of 17 (59%) doxycycline-administered *Kras*; *Gnas* mice harbored high grade precursor lesions (including mIPMNs with high-grade dysplasia in the lining epithelium), while only 6 of 18 (33%) no doxy control *Kras*^G12D^-expressing mice harbored high-grade lesions (**Figure 1G**), which is consistent with the findings observed at earlier time points (**Figure 1F**).

Despite a greater prevalence of high-grade precursor lesions, however, there was no significant difference in the frequency of invasive carcinoma upon co-expression of *Gnas*^R201C^. While the median survival of doxy administered *Kras*; *Gnas* mice were significantly shorter than control mice without doxy (**Figure 1H**). Of note, however, 8 of 10 doxy administered *Kras*; *Gnas* mice without invasive carcinoma were euthanized due to abdominal distention caused by large cyst formation within the pancreas, suggesting that measuring median survival alone was a confounding factor for assessing natural history of these mice.

In a cohort of 35 *Kras*; *Gnas* mice that was monitored for progression to invasive carcinoma (17 with doxy, and 18 without doxy administration), five mice in the doxy arm (29%) and six mice in the control arm (33%) developed invasive neoplasms at a median interval of 43 weeks. All of the invasive carcinomas developing in the doxy-induced *Kras*; *Gnas* mice were well to moderately differentiated ductal adenocarcinomas (**Figure 2A, Supplementary Table 2**); no colloid carcinomas, which are mucin-rich invasive cancers arising in the context of intestinal type human IPMNs, were observed. On the other hand, control (no doxy) mice predominantly develop poorly differentiated or undifferentiated adenocarcinomas (5 of 6 cases). Of the mice that developed cancers in each cohort, *Kras*; *Gnas* mice with doxy administration exhibited predominantly locally invasive PDAC without distant organ metastasis (4 of 5 cases), while control mice exhibited somewhat higher metastatic rate (3 of 6 cases). Immunohistochemical assessment of phospho-Map kinase (pMAPK) expression in PDAC arising in *Kras*; *Gnas* mice with or without doxy showed distinctly higher expression of pMAPK upon *Gnas*^R201C^ coexpression (**Figure 2B**), consistent with the additive effect of cAMP-signaling on activation of the MAP kinase pathway.

**Figure 2.**
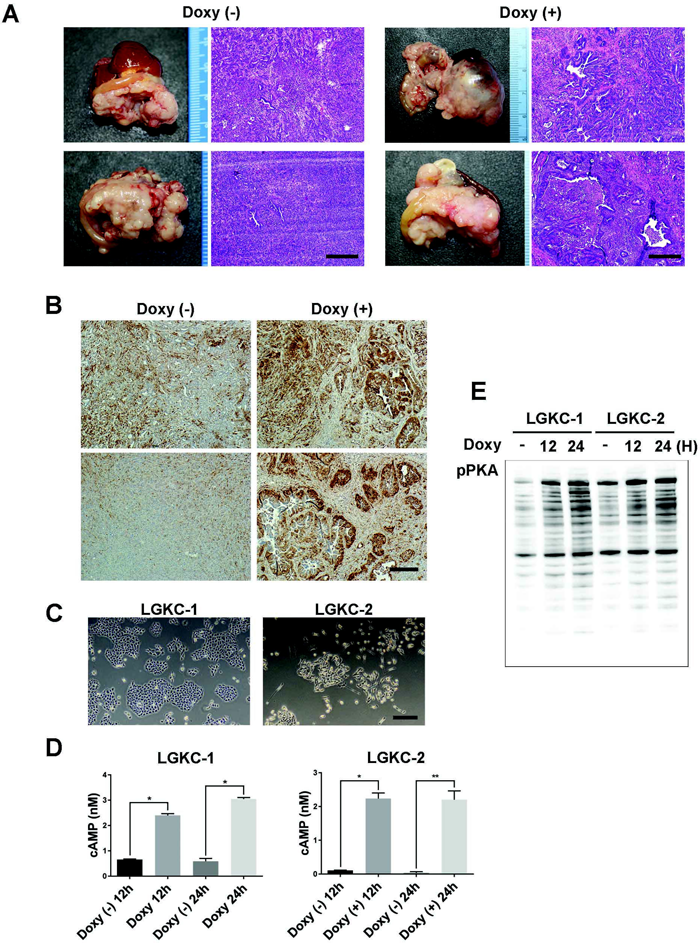
Generation of isogenic pancreatic cancer cell lines with inducible cyclic AMP (cAMP) activation from invasive carcinomas arising in *Kras*; *Gnas* mice. (A) Representative gross appearance and histology of invasive carcinomas arising in the pancreas of *Kras*; *Gnas* mice maintained on either control diet or doxycycline diet (scale bar, 50µm). (B) Immunostaining for phospho-mitogen-activated protein kinase (pMAPK) in invasive carcinomas arising in pancreata of *Kras*; *Gnas* mice fed either control diet or doxycycline diet (scale bar, 25 m). (C) Morphology of two independent isogenic cell lines – LGKC-1 and LGKC-2 – generated from invasive carcinomas arising in *Kras*; *Gnas* mice. Mutant *Kras*^G12D^ allele is constitutively expressed in the lines, while *Gnas*^R201C^ expression can be induced by addition of doxycycline (scale bar, 100µm). (D) LGKC-1 and LGKC-2 cells show significant upregulation of cAMP levels upon doxycycline induced *Gnas*^R201C^ activation at 12 and 24 hours post-induction. (E) Western blot analysis confirms activation of phospho-protein kinase A (pPKA) substrates in LGKC-1 and LGKC-2 cells upon doxycycline induced *Gnas*^R201C^ activation.

### Co-expression of *Gnas*^R201C^ on *Kras*^G12D^ background paradoxically attenuates the oncogenic phenotype of murine PDAC cells

For further mechanistic analysis, we established PDAC cell lines from primary tumors that developed in the pancreata of 2 independent *Kras*; *Gnas* mice at the age of 8 and 10 months, respectively, which we named Linker; *Gnas*; *Kras*.; Cre-1 (LGKC-1) and LGKC-2 cell lines (**Figure 2C**). Both of these cell lines represent an isogenic system, which constitutively express *Kras*^G12D^, but *Gnas*^R201C^ is only co-expressed upon addition of doxy *in vitro*. Therefore, these lines can serve as a conduit for elucidating how aberrant cAMP alters signaling pathways on the backdrop of constitutively active Ras. Indeed, *in vitro* assays confirmed that both LGKC-1 and LGKC-2 lines significantly upregulated cAMP levels upon doxy administration (**Figure 2D**), as well as phospho-Protein Kinase A (pPKA) substrates (**Figure 2E**).

We then assessed anchorage-independent growth and invasiveness of LGKC-1 and LGKC-2 lines with or without doxy administration. Rather unexpectedly, the number of colony in soft agar was significantly decreased in both LGKC-1 and LGKC-2 lines by the induced expression of *Gnas*^R201C^, with a more profound reduction of colony formation observed in the LGKC-2 line (**Figure 3A**). On the same lines, invasive capacity of both lines was also attenuated in an *in vitro* Matrigel assays, upon *Gnas*^R201C^ induction (**Figure 3B**). Mirroring the paradoxical *in vitro* data, subcutaneous allografts established from both LGKC-1 and LGKC-2 lines in doxy-fed athymic mice demonstrated reduced tumor weights compared to control allografted mice (**Figure 3C**); albeit, in contrast to LGKC-2 allografts, in LGKC-1 cells, the reduction in mean tumor weights did not reach statistical significance. Overall, however, the compendium of *in vitro* and *in vivo* data underscored the surprising finding that co-expression of *Gnas*^R201C^ on a *Kras*^G12D^ background attenuated the transformed phenotype for established PDAC cells. The most remarkable difference was however in the histology of isogenic allografted tumors, where expression of *Gnas*^R201C^ was with doxy resulted in striking epithelial differentiation, which was not apparent in matched allografts established without doxy (**Figure 3D**). This effect of mutant *Gnas*^R201C^ on PDAC differentiation seems to be consistent with the findings in autochthonous *Kras*., *Gnas* mice. The epithelial differentiation induced by *Gnas*^R201C^ was more prominent in LGKC-1 line, compared to LGKC-2, but this was not surprising given that the isogenic LGKC-2 allografts without doxy were essentially undifferentiated.

**Figure 3.**
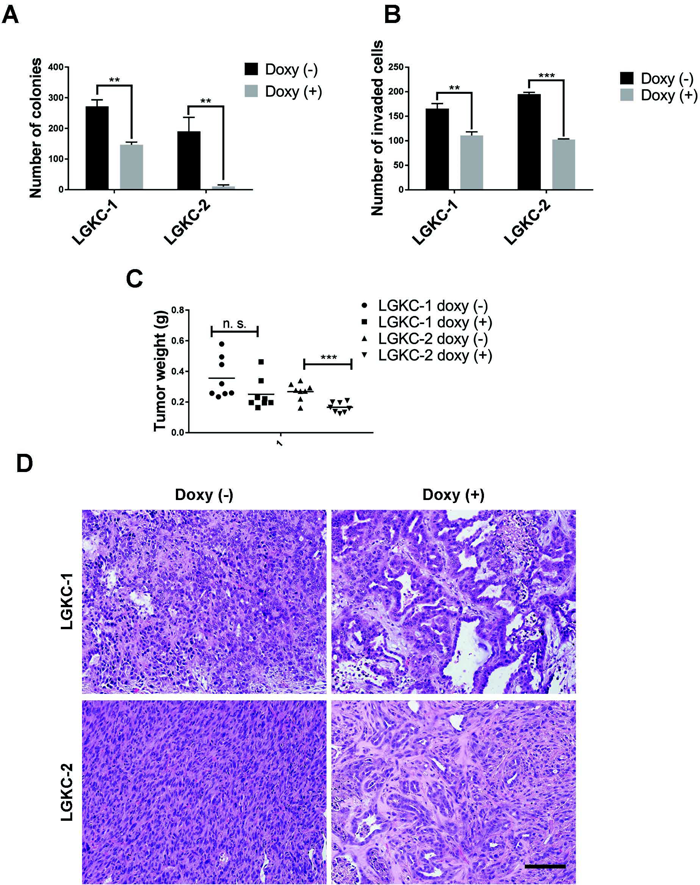
Expression of *Gnas*^R201C^ allele in LGKC cells induces a striking differentiation phenotype *in vivo*. (A) Quantification of colony formation in LGKC-1 and LGKC-2 cells treated with or without doxycycline for 2 weeks (* = *P* <0.05). (B) Quantification of invasion assays in LGKC-1 and LGKC-2 cell treated with or without doxycycline for 48 h (** = *P* <0.01). (C) Subcutaneous allografts derived from LGKC-1 (*left*) and LGKC-2 cells (*right*) in athymic mice fed either normal diet or doxycycline diet (n = 8 mice for each cell line and doxycycline treatment) (* = *P* <0.05). (D) Representative histology of LGKC-1 (*top panels*) and LGKC-2 allografts (*bottom panels*) in athymic mice fed either normal diet or doxycycline diet (scale bar, 50 m).

### Co-expression of *Gnas*^R201C^ on *Kras*^G12D^ background activates upstream Hippo kinase cascade and attenuates YAP1 signaling

We probed for the potential downstream pathways that might account for this striking induction of differentiation in mutant *Kras*^G12D^ tumors upon co-expression of *Gnas*^R201C^. Microarray analysis of the isogenic lines showed a prominent YAP1 repression signature in LGKC lines upon doxycycline induction (**Supplementary Figure 3A**), suggesting that cAMP signaling might be suppressing this critical transcriptional co-activator implicated in PDAC pathogenesis. Indeed, in both LGKC-1 and LGKC-2 cells, activation of *Gnas*^R201C^ signaling was associated with significant reduction in credentialed YAP1 target genes (m*Ctgf*, m*Cyr61*, m*Thbs1*, m*Itgb5*), albeit the effects were more striking in LGKC-2 cells (**Supplementary Figure 3B**). As a transcriptional co-activator, activated YAP1 is localized in the nucleus, while cytoplasmic sequestration is indicative of phosphorylation and inactivation by the upstream Hippo kinase cascade. Indeed, upon *Gnas*^R201C^ activation, YAP1 was substantially sequestered in the cytoplasm in both LGKC-1 and LGKC-2 cells (**Figures 4A-B**), consistent with inactivation. Assessment of nuclear and cytoplasmic extracts also confirmed evidence for cytoplasmic sequestration upon *Gnas*^R201C^ activation (**Figure 4C**), with no changes in the levels of the main YAP1 binding partner and transcription factor Tead2. In LGKC-1 and LGKC-2 cells, doxycycline induction led to time-dependent phosphorylation of the immediate upstream kinase Lats1, which in turn, phosphorylates and inactivates YAP1 at two distinct serine residues (S127 and S397) (**Figure 4D**). The impact of doxycycline induction on YAP1 activation status in both LGKC lines was pheno-copied by the cAMP elevator forskolin (**Figure 4E**), which further reiterated that the effects were through the canonical signal transduction pathway downstream of Gas. Orthogonal validation for a role of mutant *Gnas*^R201C^ activation in regulating YAP1 activation through cytoplasmic sequestration was borne out by immunohistochemical staining for YAP1 expression in *Kras;Gnas* mice, with or without doxycycline (**Figure 5A**). In contrast to the strong nuclear localization of YAP1 protein in pancreatic precursor lesions (mPanINs) of all histological grades in mice without doxycycline administration (*left*), cytoplasmic localization was the predominant pattern seen in *Kras;Gnas* mice receiving doxycycline (*right*). This suggests that the impact of *Gnas*^R201C^ activation on the Hippo pathway is an early event, occurring at the step of pre-invasive neoplasia.

**Figure 4.**
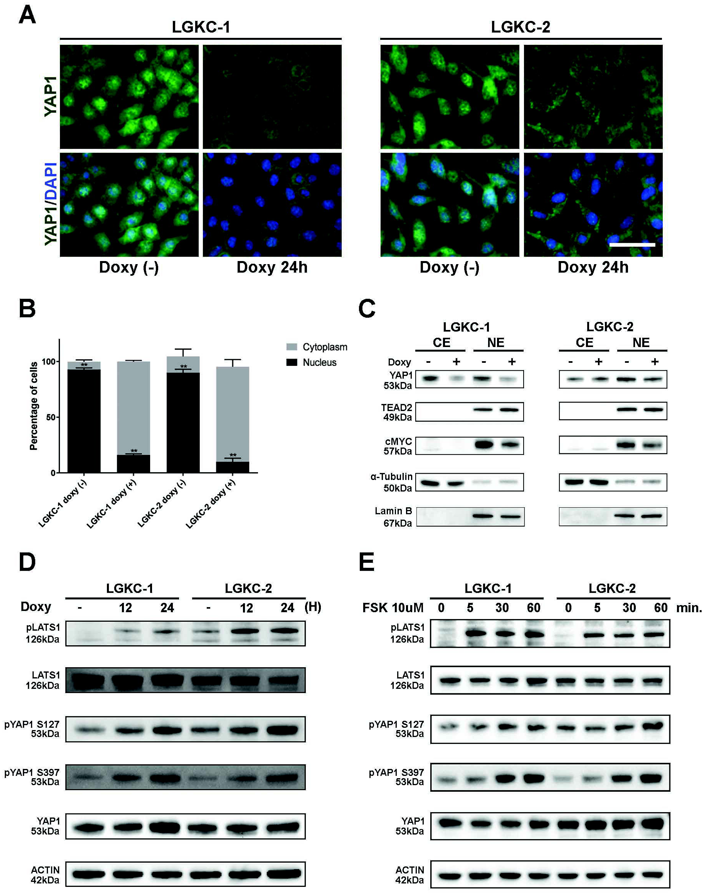
Induction of mutant *Gnas*^R201C^ leads to YAP1 cytoplasmic sequestration through the activation of Hippo tumor suppressor pathway. (A) Representative photomicrographs of non-confluent LGKC-1 (*left*) and LGKC-2 cells (*right*) untreated or treated with doxycycline for 24 hours and stained for YAP1 (green) and nuclei (DAPI, blue) (scale bar; 50µm). (B) The proportion of cells per field showing YAP1 localization in the nucleus or cytoplasm. n=3 independent experiments. (C) Nuclear and cytoplasmic extracts from LGKC-1 and LGKC-2 cell lines demonstrates reduced nuclear YAP1 localization upon doxycycline induction. Of note, c-Myc protein levels (a YAP1 target gene) are also reduced in both cell lines upon doxycycline induction. (D) Western blot analysis LGKC-1 and LGKC-2 cell lines demonstrates activation of the upstream Hippo kinase cascade (specifically phosphorylation of LATS1) upon doxycycline induction, which is accompanied by inactivating phosphorylation of YAP1 at residues S127 and S397. (E) The cAMP elevator, forskolin, phenocopies the effects of Gαs induction on LATS1 and YAP1 S127/S397 phosphorylation.

As the most profound impact of *Gnas*^R201C^-dependent YAP1 inactivation appears to be on the differentiation status of tumors, we assessed whether sustained YAP1 activity would “rescue” this phenotype. In paired allograft experiments, where we stably expressed either wild type *YAP1*, or the phosphorylation resistant *YAP1*^S127A^ mutant allele in LGKC-1 cells, the differentiation phenotype observed with doxycycline induction was completely abrogated when *YAP1*^S127A^ was expressed in LGKC-1 allografts (**Figure 5B**). To further explore the mechanistic basis of this striking epithelial differentiation phenotype, we examined the dynamics of α-E-catenin localization in LGKC cells, derivative allografts and in the autochthonous models, with and without induction of *Gnas*^R201C^ expression. In the epidermis, membranous localization of α-E-catenin has been implicated as a mediator of epidermal differentiation, wherein it also plays a role as a tumor suppressor through sequestering YAP1 in the cytoplasm ^23^. Under sub-confluent conditions, addition of doxycycline to LGKC-1 cells was associated with strong membranous expression of α-E-catenin that coincided with cytoplasmic YAP1 sequestration (**Figure 6A**). Moreover, in subcutaneous allografts generated from LGKC-1 cells, the previously described epithelial differentiation phenotype was also similarly associated with membranous α-E-catenin localization and nuclear YAP1 exclusion (**Supplementary Figure 4A**); this differential compartmentalization was also phenocopied in the autochthonous model. Specifically, while mPanIN lesions in *Kras;Gnas* mice with or without doxycycline induction expressed strong α-E-catenin membrane localization, this pattern was lost in the setting of mutant *Kras*^G12D^ induced invasive cancers, while retained in the differentiated PDAC that arose in the setting of concomitant *Gnas*^R201C^ expression (**Figure 6B**).

**Figure 5.**
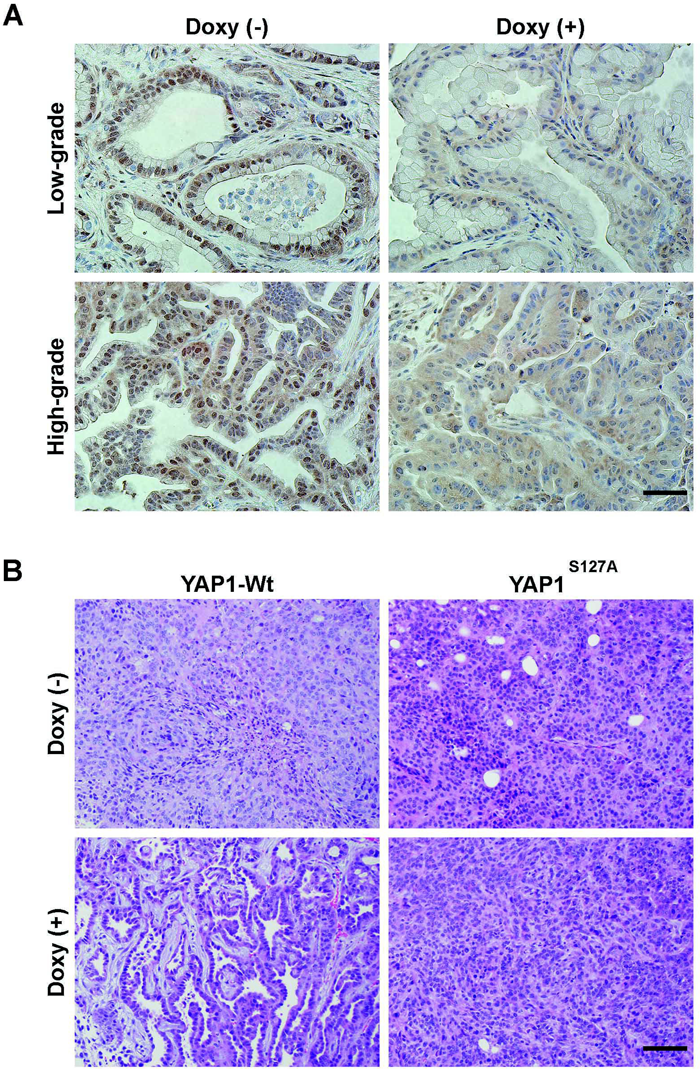
Cystic neoplasia in the murine pancreas progresses along a YAP1 independent pathway and can be bypassed using a phosphorylation resistant *YAP1* mutant allele. (A) Immunohistochemical assessment of YAP1 nuclear localization shows reduced nuclear YAP1 localization in both low-grade and high-grade pancreatic precursor lesions arising in *Kras*; *Gnas* mice fed doxycycline *versus* those on control diet (scale bar, 50µm). (B) In LGKC-1 allografts established with cells expressing the Hippo phosphorylation resistant *YAP*^S127A^ allele, expression of doxycycline-induced Gαs fails to induce the differentiation phenotype (*right panels*) observed with wild type YAP1 expression (*left panels*) (scale bar, 100µm).

**Figure 6.**
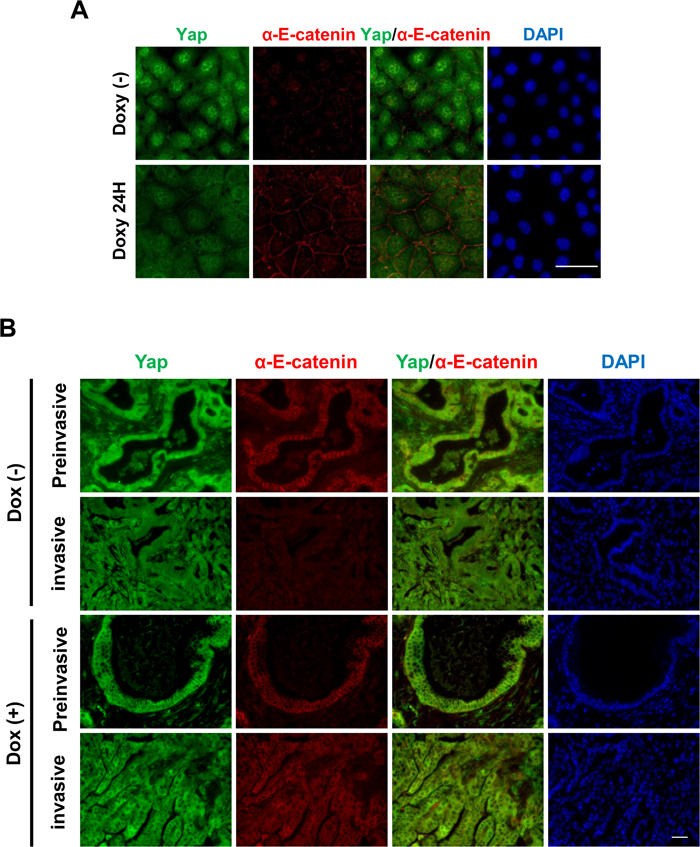
Cytoplasmic sequestration of YAP1 induced by mutant *Gnas* enhances membranous expression of α-E-catenin. (A) Representative photomicrographs of immunofluorescence assay for YAP1 and α-E-catenin localization in LGKC-1 cells treated with (*bottom panels*) or without doxycycline (*top panels*) for 24 h. Green = YAP1; Red = α-E-catenin; DAPI = nucleus, blue (scale bar, 50µm) (B) Representative photomicrographs of immunofluorescence assay for YAP1 and α-E-catenin localization in precursor lesions and invasive cancers arising in autochthonous *Kras*; *Gnas* mice with (*bottom two panels*) or without (*top two panels*) doxycycline treatment. While membranous α-E-catenin expression is retained in pancreatic precursor lesions irrespective of Gαs expression (i.e. irrespective of doxycycline treatment), only the differentiated invasive cancers arising in the doxycycline-fed, Gαs expressing *Kras*; *Gnas* mice show retained membranous α-E-catenin localization. In contrast, minimal membranous α-E-catenin localization is seen in the invasive cancers arising in the absence of Gαs expression (scale bar, 50μm).

### YAP1 localization in human IPMNs in relationship to mutant *GNAS* status

In light of the impact of *GNAS* mutation on YAP1 localization in murine tissues, we performed immunohistochemical staining for YAP1 in human IPMNs with known *GNAS* mutations (either *GNAS*^R201C^ or *GNAS*^R201H^). Seven of 8 (88%) *GNAS* wild type IPMNs exhibited strong nuclear staining for YAP1 (**Supplementary Figure 4B**, *left*); in contrast, 6 of 14 (44%) IPMNs with mutant *GNAS* had minimal YAP1 expression in IPMN lesional tissue, despite robust internal controls (ducts and centroacinar cells that expressed strong nuclear YAP1 on the same section) (*data not shown*). Of the remaining 8 of 14 *GNAS* mutant IPMNs, two (14%) had expression restricted to the cytoplasm only (see representative image in **Supplementary Figure 4B**, *right*), three (21%) had a higher score of YAP1 expression in the cytoplasm compared to the nucleus (consistent with cytoplasmic sequestration), and three (21%) had similar levels of expression in both compartments (**Supplementary Table 3**). While we did not observe an obvious effect of *GNAS* mutations on the grade of IPMN differentiation, it is noted that the 22 lesions we queried with known *GNAS* status were non-invasive IPMNs, and there were no PDAC arising from these lesions.

## Discussion

Our study demonstrated that targeted expression of a mutant *Gnas*^R201C^ allele in the adult mouse pancreas, concurrently with an activating *Kras*^G12D^ mutation, induces the development of pancreatic cystic neoplasms, including their eventual multistep progression to PDAC, which mirrors the cognate IPMN-PDAC progression model in humans. In addition, we demonstrate that canonical Gαs-cAMP-PKA signaling activates the inhibitory Hippo kinase cascade and sequesters the transcriptional co-activator protein YAP1 in the cytoplasm, which has a profound impact on the differentiation status of invasive carcinomas that arise in the context of mutant *Gnas*^R201C^. Although a previous study has reported a mouse model displaying IPMN-like pancreatic cystic lesions by the co-expression of mutant *Kras*^G12D^ and *Gnas*^R201C^ alleles during embryonic development, this model did not recapitulate the entire spectrum of IPMN progression because of the early death of mice (median, 5 weeks) ^24^. Thus, our genetically engineered mouse (GEM) model of IPMNs provides the first *bona fide* cross-species platform where we can potentially generate longitudinal data points for biomarker and imaging studies, with the intent of translational research in early diagnosis and interception of PDAC

Pancreas-specific expression of mutant *Tp53*, or inactivation of *Smad4* or *Cdkn2A*, demonstrate robust oncogenic cooperation with mutant *Kras*^G12D^, leading to accelerated formation of mPanINs, and invasive carcinoma with higher frequency (ranging from 38 to 96%) and shorter latency (median 14 to 25 weeks) ^25–27^, respectively, in comparison to mice with pancreas-specific *Kras*^G12D^ mutation alone ^28^. On the contrary, in the *Kras;Gnas* mice, we did not observe the stated acceleration, with a median time to invasive carcinoma of 46 weeks (38 weeks post doxycycline) comparable to control mice, suggesting that *Gnas*^R201C^ is not functioning as a traditional cooperating oncogene, but rather, as an “onco-modulator” that alters the phenotype of pre-invasive lesions arising in the context of mutant *Kras*^G12D,^ and leads to invasive carcinomas with a predominantly well differentiated morphology. Expression profiling studies identified repression of the transcriptional co-activator YAP1 as a signature of *Gnas*^R201C^ co-expression on the backdrop of mutant *Kras*^G12D^, through activation of the upstream inhibitory Hippo kinase cascade. Notably, YAP1 is inactivated through phosphorylation-dependent cytoplasmic sequestration, which we observed not only in murine autochthonous or allograft models upon induction of Gαs, but also in patient-derived IPMNs specimens with known *GNAS* mutations. We were able to demonstrate the primacy of YAP1 sequestration in the Gαs-induced differentiation phenomenon, and the role of Hippo-induced phosphorylation as the mediator of this effect, through “rescue” experiments performed with a phosphorylation resistant mutant allele that completely reversed the differentiation effect *in vivo*.

YAP1 has been implicated as a critical transcriptional co-activator in both development and neoplasia, including that of the pancreas. We, and others, have previously demonstrated the transforming capabilities of *YAP1* as an oncogene in the pancreatic epithelium ^29, 30^; specifically, YAP1 has been demonstrated to exhibit both cancer cell intrinsic growth properties (through induction of JAK-STAT3 and MYC signaling, amongst others) ^31, 32^, as well as a profound impact on the tumor microenvironment by creating a permissive immune milieu ^33^. Conversely, genetic ablation of *YAP1* in the murine pancreas on a mutant *Kras* or *Kras;p53* background blocks progression to PDAC, although the earliest precursor lesions (acinar ductal metaplasia) are not impeded ^34^. Notably, despite the abrogation of YAP1 signaling in our model, we do encounter eventual progression in a subset of mice to invasive cancers, suggesting that constitutive cAMP activation likely has additional downstream effects that sustain the neoplastic phenotype. Another key difference from the prior study pertains to embryonic ablation of YAP1 ^34^, which likely has more profound impact on the pool of pancreatic progenitors amenable to neoplastic transformation ^35^, than Gαs-induced inhibition of YAP1 in the adult pancreas. Irrespective, we believe that the inhibition of YAP1 signaling plays a profound role in attenuating the unchecked progression of IPMNs to PDAC, and could explain the favorable biology of these cystic neoplasms. This recalcitrance to invasive neoplasia might also explain why *GNAS* mutations are uncommon in PanIN lesions, which are the more common precursor to “conventional” adenocarcinoma and harbor a distinct natural history compared to IPMN-induced cancers.

While phosphorylation of YAP1 through the Hippo kinase cascade results in cytoplasmic sequestration, there is a strong selection pressure during carcinogenesis to unleash this important oncogenic signal through phosphatase-dependent de-phosphorylation and nuclear translocation. Prior studies in epidermal keratinocytes and breast epithelium have shown the importance of the cell-cell adhesion protein α-E-catenin in “fencing in” phosphorylated YAP1 ^23, 36–38^, and shielding the phosphorylated protein from phosphatases like PP2A. Further, downregulation of α-E-catenin in cutaneous squamous cell carcinoma models was shown to result in poorly differentiated cancers with nuclear YAP1 translocation ^23, 36^. In *Kras;Gnas* mice, we find that Gαs-induced YAP1 sequestration in the cytoplasm is accompanied by robust expression of membranous α-E-catenin, likely functioning as a cytoplasmic “restraint”, while nuclear translocation of YAP1 (seen with mutant *Kras*^G12D^ expression alone) is accompanied by loss of membrane expression of α-E-catenin. While some studies have suggested that α-E-catenin itself regulates the activity of YAP1 through a non-canonical Hippo-independent pathway ^39^, we do not find definitive evidence to this effect in our model, since Gαs induction phosphorylates the canonical Hippo kinase LATS1, and at least in the precursor lesions arising in the setting of *Kras*^G12D^ alone, we observe nuclear YAP1 expression despite robust membranous α-E-catenin localization (the latter, however, is lost in the invasive carcinomas).

In summary, we describe a murine model of IPMNs that recapitulates the genetic landscape of the most common drivers and the multistep progression observed in the cognate human cystic neoplasms. We identify the potential mechanistic basis for the relatively indolent biology of the cystic pathway to invasive neoplasia in the pancreas, and the role of constitutively active Gαs as an “onco-modulator” of epithelial differentiation in the context of mutant *Kras*. The *Kras;Gnas* model (and derivative reagents, such as isogenic cell lines) will be useful reagents for the community to pursue cross-species translational research endeavors in the context of early detection and cancer interception.

## Acknowledgements

AM is supported by the MD Anderson Pancreatic Cancer Moonshot, Cancer Prevention and Research Institute of Texas (CPRIT), NCI CA196403, NCI CA200468, NCI CA218004, and the Khalifa Bin Zayed Al Nahyan Foundation. LW is supported by the Sol Goldman Pancreatic Cancer Research Center. NI and HY were supported in part by the Uehara Memorial Foundation.

## Author contributions

NI and HY were equally contributed to this work. Study concept and design: NI, HY, JSG, AM. Data acquisition and analysis: NI, HY, TO. Material support: CF, LW, AS. Drafting of manuscript: NI, AM. Final approval of submission: NI

